# Sca1^+^ cells as direct isolate (*ex vivo*) versus *in vitro* cultured exhibit differential proteomic signatures in murine skeletal muscle

**DOI:** 10.1101/2020.12.02.409110

**Authors:** Saketh Kapoor, Pratigya Subba, Sudheer Shenoy P, Bipasha Bose

**Affiliations:** Stem Cells and Regenerative Medicine Centre, Yenepoya Research Centre, Yenepoya (Deemed to be University), Deralakatte, Mangalore, Karnataka, Pincode-575018, India; Center for Systems Biology and Molecular Medicine, Yenepoya Research Centre, Yenepoya (Deemed to be University), Deralakatte, Mangalore, Karnataka, Pincode-575018, India

**Author notes:** **Correspondence**, Bipasha Bose. Sudheer Shenoy P. **Address:** Stem Cells and Regenerative Medicine Centre, Yenepoya Research Centre, Yenepoya University, University Road, Deralakatte, Mangalore, 575 018, Karnataka, India.

**Keywords:** Stem Cell Antigen-1 (Sca-1), regenerative stem cells, stem cells proteomics, mass spectrometry-based proteomics

## Abstract

Stem cell antigen-1 (Sca-1) is a glycosyl-phosphatidylinositol-anchored membrane protein that is expressed in a sub-population of muscle stem and progenitor cell types. Reportedly, Sca-1 regulates the myogenic property of myoblasts and *Sca-1^-/-^* mice exhibited defective muscle regeneration. Although the role of Sca-1 in muscle development and maintenance is well-acknowledged, molecular composition of muscle derived Sca-1^+^ cells is not characterized. Here, we applied a high-resolution mass spectrometry-based workflow to characterize the proteomic landscape of mouse hindlimb skeletal muscle derived Sca-1^+^ cells. Furthermore, we characterized the impact of the cellular microenvironments on the proteomes of Sca-1^+^ cells. The proteome component of freshly isolated (*ex vivo*) Sca-1^+^ cells was compared with that of Sca-1^+^ cells expanded in cell culture (*in vitro*). The analysis revealed significant differences in the protein abundances in the two conditions reflective of their functional variations. The identified proteins were enriched in various biological pathways. Notably, we identified proteins related to myotube differentiation, myotube cell development and myoblast fusion. We also identified a panel of cell surface marker proteins that can be leveraged in future to enrich Sca-1^+^ cells using combinatorial strategies. Comparative analysis implicated the activation of various pathways leading to increased protein synthesis under *in vitro* condition. We report here the most comprehensive proteome map of Sca-1^+^ cells that provides insights into the molecular networks operative in Sca-1^+^ cells. Importantly, through our work we generated the proteomic blueprint of protein abundances significantly altered in Sca-1^+^ cells under *ex vivo* and *in vitro* conditions.

## Introduction

The stem cell antigen-1 (Sca-1), a member of *Ly6* gene family, is an 18-KDa glycosyl phosphatidylinositol-anchored protein that is localized in the lipid rafts of plasma membrane [1]. It is one of the most common markers of mouse hematopoietic stem cells (HSCs) [2].The expression of Sca-1 has also been identified in a variety of stem/progenitor cells from various tissues and organs [3]. In muscles, Sca-1 has been used as marker for mouse muscle-derived stem cells (MDSCs) [4] and it’s expression was also reported on the myogenic precursor cells [5]. The expression of Sca-1 has been reported on the muscle satellite cells and fibro/adipogenic progenitors which participate in muscle regeneration [6,7]. However, satellite cells remain heterogenous for the expression of Sca-1 antigen [8]. MDSCs isolated from the dystrophic *mdx* mice were shown to express Sca-1 along with early myogenic progenitor [4]. These Sca-1^+^ MDSCs when injected into *mdx* mice, helped in the regeneration of new dystrophin positive muscle fibres [4]. Further, Sca-1^+^ cells isolated from muscles of new-born mice enabled regeneration of muscle fibers when injected into *mdx* mice, leading to speculations on their role in therapeutic intervention in myopathies [9]. Additionally, cardiac Sca-1^+^ myogenic precursor cells were associated with functions in cardiac repair [10]. Sca-1 was demonstrated to regulate self-renewal of mesenchymal progenitors and the differentiation of osteoclasts [11]. Mice deficient for the expression of Sca-1^(-/-)^ demonstrated defects in muscle regeneration as well as age dependent loss of bone mass resulting in brittle bone [11,12]. In C2C12 cells, downregulation of Sca-1 expression prevented myoblast differentiation and led to defects in myotube formation [13]. Sca-1^+^ cells represent a low percentage of muscle tissue stem/progenitor cells and can be isolated using fluorescence-activated cell sorting (FACS). Despite the therapeutic value, the molecular composition of Sca-1^+^cells are far from complete.

High-throughput mass spectrometry has emerged as an efficient technique to dissect molecular mechanisms operating in biological systems [14,15]. Recently, a combination of FACS and data-independent acquisition mass spectrometry was used to generate the proteome of rare population of human hematopoietic stem/multipotent progenitor cells [16]. In muscle biology, proteomics has been used for the comprehensive characterization of muscle proteome, however, it remains challenging due the dynamic range of proteome in skeletal muscle [17]. Nevertheless, an in-depth proteomic characterization of C2C12 mouse myoblast cells and triceps muscles of C57BL/6 mice provided a detailed map of muscle proteins involved in the glucose and lipid metabolic pathways [18]. Differential regulation of molecules such as glucose transporter (GLUT), TBC1D, a regulator of GLUT and Rabs in insulin signalling and AMPK mediated glucose uptake were observed in the two samples [18]. Through another proteomics study in mice, exercise training and diet dependent expression of MUP1 protein was identified as a factor for improving insulin sensitivity in muscle cells [19]. Proteomic analysis of human skeletal muscle biopsies from individuals undergoing different levels of physical activity revealed protein levels opposite to those reported in skeletal muscle aging, emphasizing the benefits of physical activity [20]. Comparative proteomics analysis of mouse pluripotent ES-D3 cell line differentiated into Sca-1^+^ cells was also reported using two-dimensional gel electrophoresis method [21].

Currently, to the best of our knowledge there are no reports on any high-resolution mass spectrometry-based proteomics analysis of mouse muscle Sca-1^+^ adult stem/progenitor cells. In this study, we report an in-depth proteomic characterization of mouse primary Sca-1^+^ cells isolated from the hindlimb skeletal muscles. Further, a comparative analysis of Sca-1^+^ *ex vivo* and *in vitro* expanded cells was carried out using label-free quantitative proteomics. Our data serves as a molecular resource for stem cell biologists. In future, our data can be utilized for screening alternative markers for the specific isolation of muscle stem cells.

## Material and Methods

### Use of animals

This study was approved by the Institutional Animal Ethics Committee, Yenepoya (Deemed to be University). BALB/c strain of mice (Adita Bio Sys Pvt Ltd, Bangalore 1868/PO/Bt/S/16/CPCSEA) of age five weeks were used in this study. All the animals were maintained on a 12:12-hour light-dark cycle housed in a Specific Pathogen Free (SPF) facility under individually ventilated caging systems (Lab Products Inc, USA). Animals were provided standard chow diet. Mice were anaesthetized using ketamine (90 mg/kg body weight) approved as a standard humane method for sacrifice.

### Dissection of hindlimb muscle and preparation of single cell suspension

Five weeks old male BALB/c mice (n = 3) were used. Briefly, the mice were sacrificed and skeletal muscles were dissected from the hind limbs. The muscle tissue was then washed in ice cold PBS, minced with fine scissors and digested in 0.2% collagenase type 1A (GIBCO) at 37°C for 40-60 min as described earlier [22]. The muscle digests were then passed through a 70 μm cell strainer (Sartorius), followed by centrifugation at 3,000 RPM for 10 min at 4°C. The cell pellet was resuspended in RBC lysis buffer (Gibco) for RBCs depletion and incubated on ice for 10 min. Finally, the cell pellet was centrifuged and the cell pellet was processed for FACS staining.

### Flow sorting of Sca-1^+^ stem/progenitor cells

Single cell suspension was prepared from hindlimb skeletal muscle as discussed above (all the procedures were performed at 4°C unless otherwise specified). The single cell suspension containing 5.0 million cells were stained using anti-mouse Sca-1 antibody conjugated with AF488 (Thermo Fisher Scientific) and incubated for 1h. The cells were washed twice with staining buffer and resuspended in 1 mL of staining buffer. The cells were sorted based on the selection of negative and positive sort mode using Bio-Rad S3e cell sorter (Bio Rad). The results were analyzed using Prosort software (Bio Rad). The positive and negative populations were collected in muscle derived stem cell (MDSC) media. Gatings were carried out using unstained and FMO controls in which 0.2 × 10^6^ cells each were taken respectively. The sorted Sca-1^+^ cells were divided into two sets; the first set labeled as the *ex vivo* Sca-1^+^ sample from which the protein was isolated by addition of lysis buffer to the cell pellet. The second set of Sca-1^+^ cells were cultured in MDSC media for their expansion which represent the *in vitro* sample.

### Culturing of flow sorted Sca-1^+^ stem/progenitor cells *in vitro*

The second set of flow-sorted, Sca-1^+^ cells were immediately cultured in matrigel coated 60 mm dish in MDSC media containing fetal bovine serum (FBS) – 10%,glutamax – 1%, penicillin & streptomycin – 1%, non-essential amino acids (NEAA) – 1%, sodium pyruvate – 1%, β Mercaptoethanol – 0.1% and Dulbecco’s modified eagle medium (DMEM) (all from GIBCO). The cells were then incubated at 37°C at 5% CO_2_ under humidified conditions. The cells were then allowed to attain 70-80% confluency and utilized for all downstream experiments.

### Protein extraction and estimation

Proteins were extracted from the two sample types: (i) Flow sorted Sca-1^+^ cells - direct (*ex vivo*) and (ii) Flow sorted Sca-1^+^ cells expanded in cell culture (*in vitro*). The cells were lysed using 100 μL lysis buffer [4% SDS, β-mercaptoethanol (1%), 1mM EDTA (pH 7.5), 1 mM sodium orthovanadate, 2.5 mM sodium pyrophosphate, 1 mM β-glycerophosphate] (Sigma-Aldrich). Samples were sonicated on ice for 20 min followed by protein denaturation at 95°C for 10 min (Qsonica). The samples were then centrifuged at 12,000 RPM for 15 min at 4°C. Clear supernatants were aliquoted and transferred to fresh microcentrifuge tubes and stored at −80°C until further use. For estimating the proteins, one aliquot of each of the sample types were thawed on ice and protein amounts were quantified using bicinchoninic acid assay (BCA) kit (Pierce, Waltham, MA). For each sample, 100 μg protein were processed for trypsin digestion.

### Trypsin digestion and peptide fractionation

Proteins from each sample were subjected to reduction using 5 mM DTT (Sigma-Aldrich) at 60°C for 20 min followed by alkylation using 10 mM iodoacetamide (Sigma-Aldrich) for 10 min at room temperature in the dark. Proteins were precipitated using five volumes of chilled acetone (Merck) at −20°C overnight. Protein pellets were retrieved through centrifugation at 13,000 RPM for 15 min at 4°C. Acetone was discarded and protein pellets were air dried for 5-10 min and then resuspended in 100 μL of 50 mM TEABC (Sigma-Aldrich) Overnight enzymatic digestion was carried out using sequencing grade trypsin (Promega) at a ratio of 1:20 enzyme:protein at 37°C. The digested peptides were dried using a speedvac concentrator (Thermo Fisher Scientific). The dried peptides were reconstituted in 60 μL of 0.1% trifluroacetic acid (TFA) (Sigma-Aldrich). Peptides were then fractionated using SCX StageTip method as described earlier [23]. Briefly, the SCX plugs were first activated using 100% acetonitrile (ACN) (Sigma-Aldrich). followed by equilibration using 0.1% TFA. Peptide samples were bound to the column followed by two washes using 0.2% TFA. Peptides were eluted in varying concentrations of ammonium acetate (Merck). The eluted peptides were dried using speedvac concentrator.

### Mass spectrometric analysis

Mass spectrometric analysis was carried out using an Orbitrap Fusion Tribrid mass spectrometer (Thermo Fisher Scientific) coupled to Easy-nLC1200 nano-flow UHPLC (Thermo Fisher Scientific). The peptides were reconstituted in 0.1% formic acid (Sigma-Aldrich) and quantified using peptide assay kit. 1 μg peptides from each fraction was loaded onto trap column nanoViper 2 cm (3 μm C18 Aq) (Thermo Fisher Scientific). Peptide separation was carried out using EASY-Spray C18 analytical column (15 cm, 75 μm PepMap C18, 2 μm C18 Aq) (Thermo Fisher Scientific) set at 40°C. The solvent gradients used for the peptide separation were as follows: linear gradient of 2-45% solvent B (80% acetonitrile in 0.1% formic acid) over 105 minutes with a total run time of 120 mins. The flowrate was set to 5 μl/min. The nanoESI source was used to generate positively charged precursor ions. Data were acquired using a data-dependent acquisition method wherein MS1 survey scans were carried out in 375-1700 m/z range. For further peptide fragmentation using MS/MS, the most intense precursor ions were selected at top speed data dependent mode with maximum cycle time of 3 sec. Peptide charge state was set to 2-6 and dynamic exclusion was set to 40 sec along with an exclusion width of ± 20 ppm. Internal calibration was carried out using lock mass option (m/z 445.12003) from ambient air. Data for the *ex vivo* and *in vitro* sets were acquired in triplicates and in total 18 raw files (2 sample types × 3 SCX fractions × 3 replicates) were generated.

### Protein identification and data analysis

The data acquired on Orbitrap Fusion Tribrid mass spectrometer (Thermo Fisher Scientific) were processed to generate peak list files using Proteome Discoverer software version 2.2 (Thermo Fisher Scientific, Bremen, Germany). The data were searched against the *Mus musculus* proteome database downloaded from NCBI FTP. The search parameters included trypsin as the proteolytic enzyme with maximum of one missed cleavage allowed. Oxidation of methionine and acetylation of protein at the N-terminus were set as a dynamic modification while carbamidomethylation of cysteine was set as static modification. Precursor and fragment mass tolerance were set to 10 ppm and 0.05 Da, respectively. A false discovery rate (FDR) value of 1% was set for the identification at protein, peptides and PSM level. For quantification of peptides across the *ex vivo* and *in vitro* datasets, the Minora Feature Detector node was activated for performing label free quantification (LFQ) workflow in Proteome Discoverer version 2.2.

### Bioinformatics analysis

The bioinformatic analysis was performed using Perseus software (http://www.perseusframework.org/) and R studio tool. Gene Ontology (GO) enrichment analysis was performed using bio-conductor package along with library (clusterProfiler) and library (org.Mm.eg.db) in R studio tool. Pathway analysis was enriched using KEEG pathways in R studio tool with the bio-conductor package. GOChord plot was generated to link the pathways to their respective genes using R studio tool [24]. GOCircle plot was created to visualize the significant pathways using R studio tool [24].

### Statistical analysis

The reproducibility of the data was analyzed by estimating the Pearson’s correlation coefficient between the technical replicates. Coefficient of Variation (CV) was calculated to determine the variation in the intensity values between the technical replicates. Fold change was calculated between proteins that were commonly identified between the Sca-1^+^*ex vivo* and *in vitro* conditions and statistical analysis was performed using Student’s t-test.

### Data availability

The MS raw files (.raw) and Proteome Discoverer search files(.msf) are available at http://www.ebi.ac.uk/pride/archive/ with the PRIDE dataset identifier PXD022247. The curated data can also be visualized at https://yenepoya.res.in/database/Sca-1-Proteomics.

## Results

### Flow sorting of mouse hindlimb skeletal muscle digests

Sca-1 plays a vital role in the development and regeneration of skeletal muscles. However, the precise mechanism through which Sca-1 is regulated and molecular networks through which Sca-1 mediates its action is poorly understood. Using the flow-sorting method, we isolated Sca-1^+^ cells from mouse hindlimb muscle tissues (Fig. 1). The percentage of pure population of Sca-1^+^ cells was 54.14% indicating the presence of a large population of Sca-1^+^ cells in the hindlimb of mouse skeletal muscle. Processing of the pure populations of Sca-1^+^ cells into two sets as *ex vivo* and *in vitro* has been mentioned in the graphical abstract.

**Figure 1:**
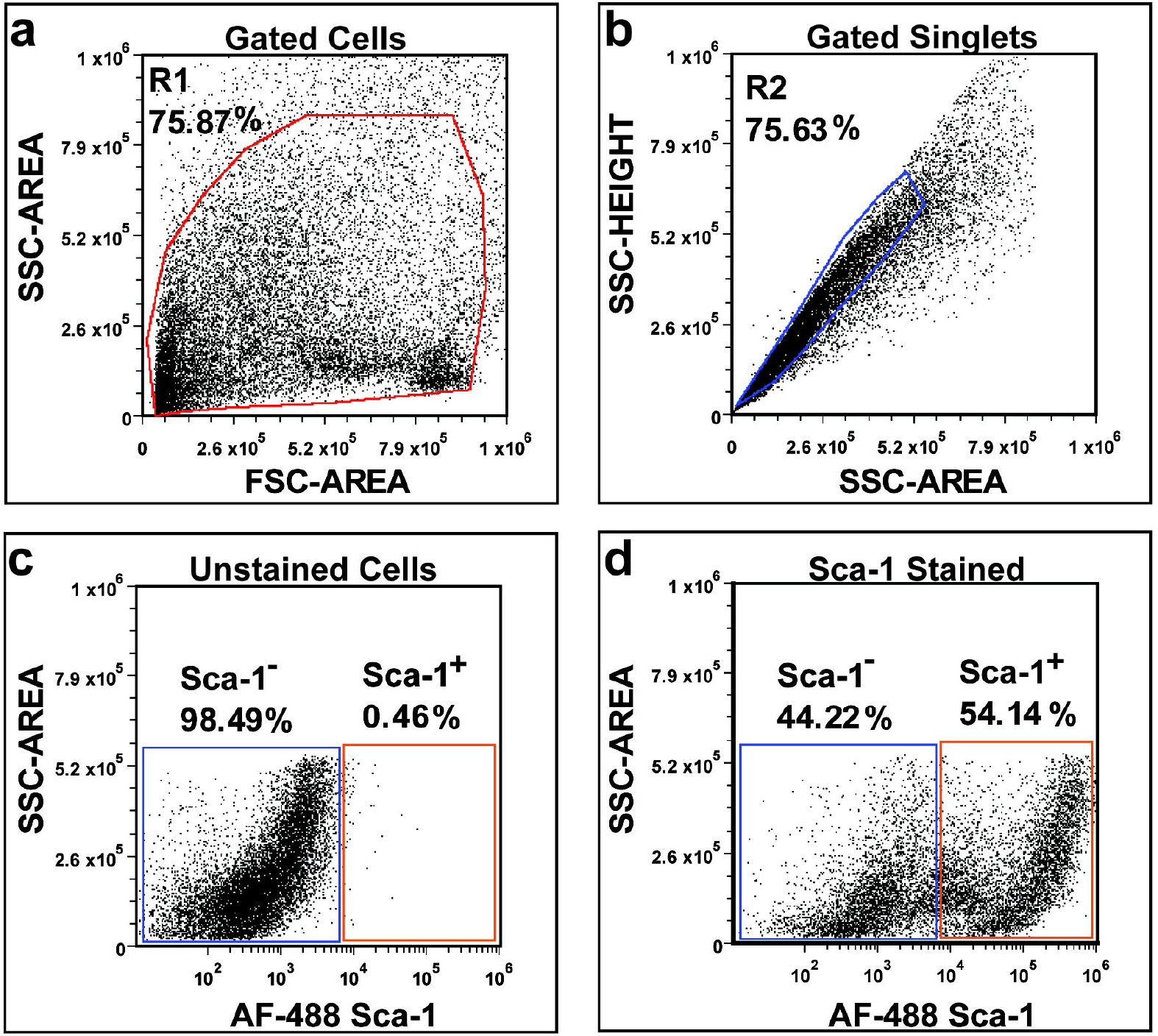
Flow sorting of Sca-1^+^ cells from skeletal muscle of five weeks old BALB/c mice. **(a)** Scatter plot depicting the cell size represented by forward scatter (FSC-Area) and cell granularity represented by side scatter (SSC-Area) gated as R1. **(b)** The singlet population gated as R2 to determine the doublet discrimination. **(c)** Unstained cells showing the respective gates to mark the positive and negative population. **(d)** Sca-1 stained cells showing the Sca-1^-^ as well as Sca-1^+^ population. The percentage are shown in respective plots.

### Proteome atlas of Sca-1^+^ muscle cells

A high-resolution proteome map of Sca-1^+^ cells was generated using an Orbitrap Fusion Tribrid mass spectrometer. The combined data of *ex vivo* and *in vitro* sets consisted of 1,109,437 MS/MS spectra and 281,373 PSMs. With a false discovery rate (FDR) set to 1% at the peptide and protein levels, 2,775 protein groups (referred to as proteins here) were identified in the *ex vivo* dataset and 4,882 proteins were identified in the *in vitro* dataset (Fig. 2a). While 2076 proteins were common to both *ex vivo* and *in vitro*, 699 proteins were unique to *ex vivo* and 2806 proteins were unique to the *in vitro* set. The complete list of proteins identified in each set and their corresponding details are provided in Supplementary Table S1. The Pearson’s correlation coefficient was equal to 1 in case of intra-sample technical replicates and the correlation coefficients dropped to ~0.45 among the inter-sample comparisons (Fig. 2b). This indicated a strong correlation of data between the technical replicates of the same sets while maintaining low correlation between the *ex vivo* and *in vitro* datasets. As indicated by the Principal component analysis, the data from technical replicates of each set clustered together tightly whereas the *ex vivo* and *in vitro* data demonstrated clear segregation (Fig. 2c). Peptide quantification across the triplicate runs was benchmarked by assessing the percentage of coefficient of variation (CV). Very high reproducibility in peptide quantification was observed with over 91% of proteins in *ex vivo* and 92% proteins in *in vitro* sets displaying CV below 20% (Fig. 2d, e). The dynamic range of proteins identified in the *ex vivo* and *in vitro* datasets spanned over several orders of magnitude (Fig. 3a, b). Low abundant proteins such as transcription factor Stat3 and kinase Akt were quantified with high intensity in both the datasets alongside highly abundant proteins such as actin and titin.

**Figure 2:**
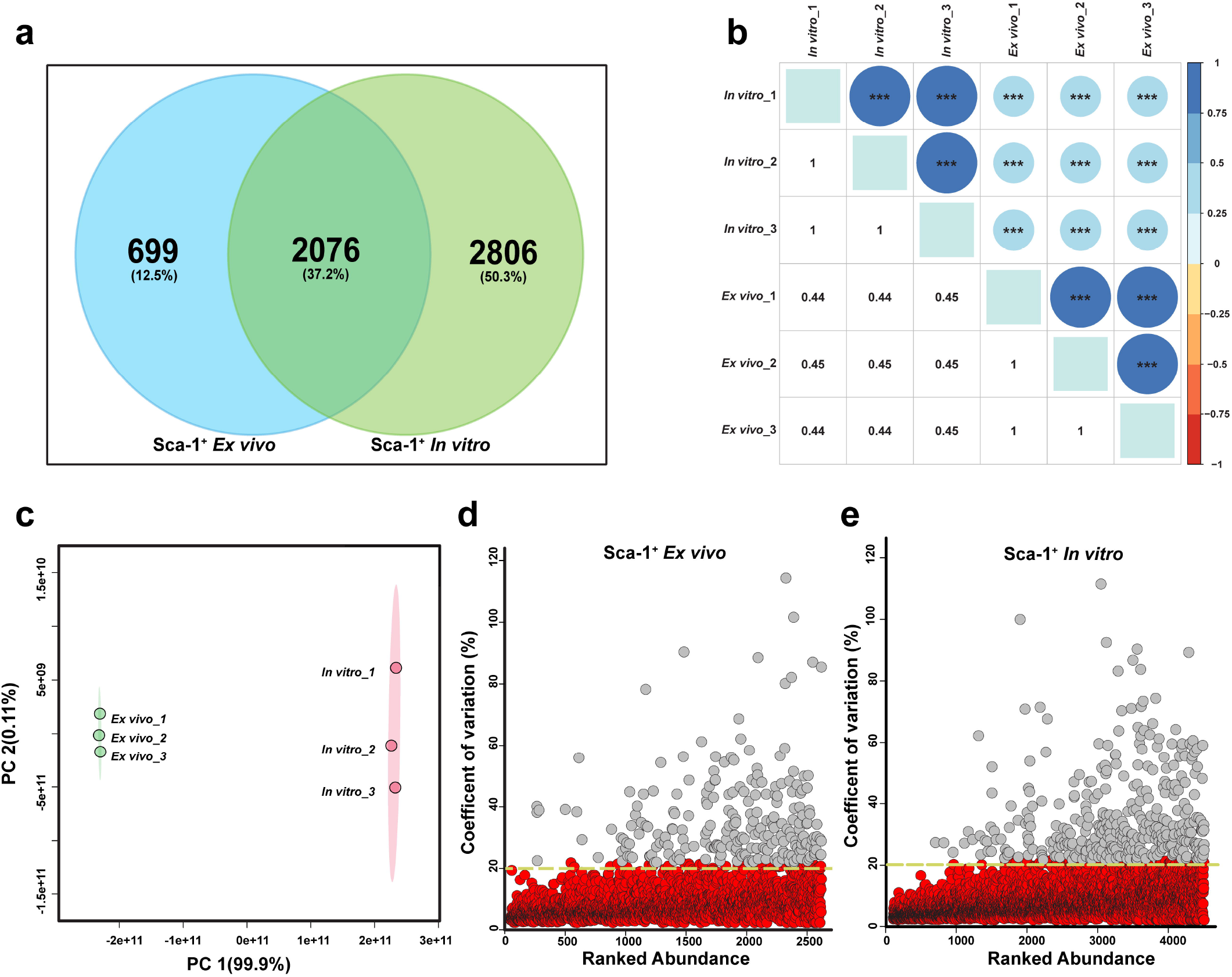
Proteomic analysis of Sca-1^+^ *ex vivo* and *in vitro* cells. (**a**) Venn diagram showing the total number of proteins identified in Sca-1^+^ *ex vivo* and *in vitro* cells. (**b**) Correlation plot show a high degree of correlation between the technical replicates in each condition. The Pearson correlation coefficient value is indicated and the p-value <0.05 (***) is considered significant. (**C**) Principal component analysis (PCA) to visualize the variations between the technical replicates of Sca-1^+^ *ex vivo* and Sca-1^+^ *in vitro* cells. **(d & e)** Coefficient of variation percentage (CV%) was assessed between the technical replicates in each run. Proteins with CV% less than 20% are marked in red and those with CV% more than 20% are marked in grey.

### Functional annotation reveals enrichment of regulatory proteins

Functional annotation was assigned to proteins identified in *ex vivo* and *in vitro* datasets based on three gene ontology (GO) terms viz., molecular function, biological process and cellular component (Fig. 3c, d). The most enriched class of proteins based on the biological process was ‘mRNA processing’ in the *ex vivo* set and ‘ribonucleoprotein binding complex’ in the *in vitro* set. Several terms such as ‘muscle contraction, ‘muscle cell development’ and ‘muscle cell differentiation’ were significantly enriched in the *ex vivo* set. However, the terms enriched in the *in vitro* set predominantly belong to ‘regulation of translation’, ‘translational initiation’ and ‘muscle cell differentiation’. Among the molecular functions, ‘actin-binding’ was most enriched in the *ex vivo* set whereas ‘mRNA binding’ was enriched in the *in vitro* set. Based on the subcellular localization, proteins of the *ex vivo* and *in vitro* sets originated predominantly from cell structures including ‘myelin sheath’, ‘actin cytoskeleton’ and ‘mitochondrion’ (Fig. 3c, d). Transcription factors are key determinants of muscle development and regeneration [25–27]. Expression of transcription factors such as Myod1, Myf5 and Mef2 are known to regulate the expression of muscle-specific genes. We investigated the representation of transcription factors by comparing our dataset with the list of mouse transcription factors in the Animal-TFDB 3.0 database. In total, we identified 170 transcription factors including MyoD1, Foxo1 and Stat3 whose roles are well described in muscle development (Supplementary Table S2). We also queried the representation of protein kinases in our dataset by comparing our data against the ‘The Mouse Kinome’ database. We identified 113 kinases that include Akt1, Akt2 and Mtor that perform distinct roles in muscle differentiation [28] (Supplementary Table S2). We also identified 34 phosphatases in our dataset when we compared our data against the data curated on mouse phosphatases [29] (Supplementary Table S2). The identification of a large number of regulatory proteins illustrates an in-depth profiling encompassing low abundant proteins.

**Figure 3:**
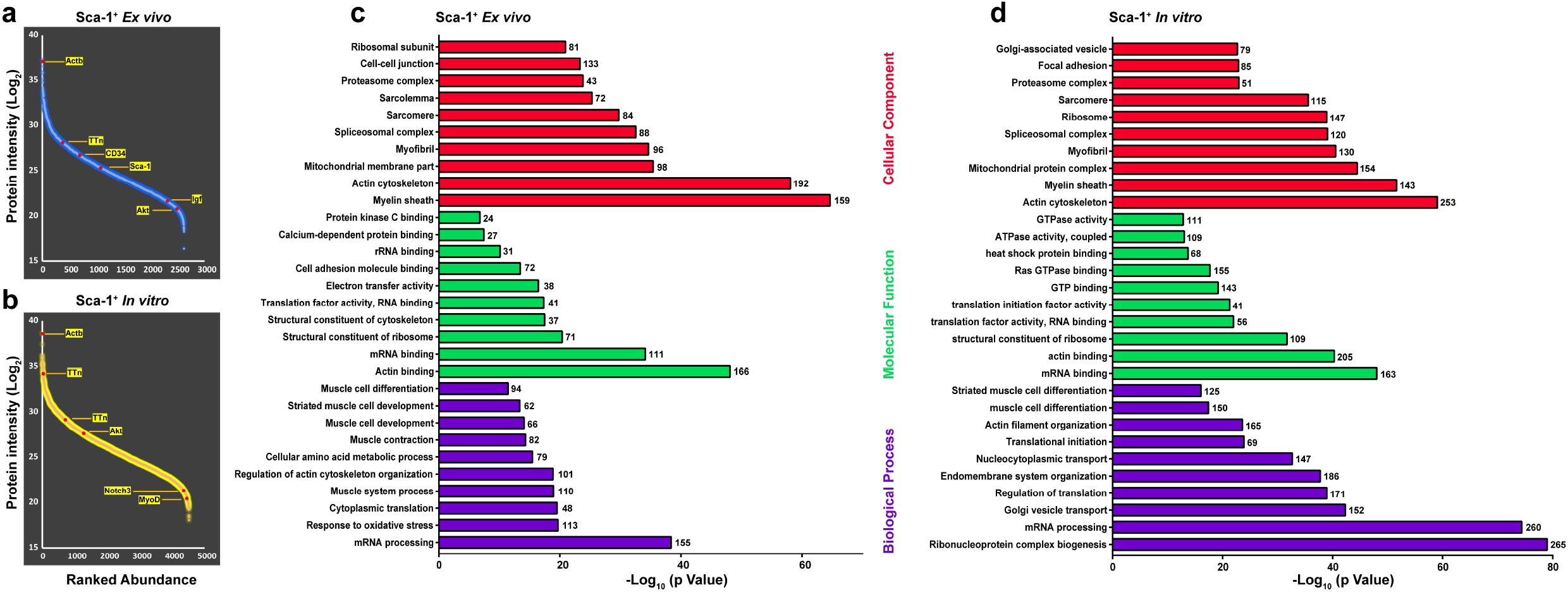
Gene-ontology of proteins identified in Sca-1^+^ dataset. Proteins are ranked based upon their abundance from highest to the lowest in Sca-1^+^ *ex vivo* (**a**) and Sca-1^+^ *in vitro* (**b**). Exemplary proteins are highlighted in each condition. Gene ontology analysis based on cellular component, molecular function and biological process in Sca-1^+^ *ex vivo* (**c**) and Sca-1^+^ *in vitro* (**d**). Those significantly associated (p ≤ 0.05) with the gene list are plotted with the numbers for each term.

### Comparative proteome analysis of *ex vivo* and *in vitro* Sca-1^+^ muscle cells

As cells for this study were sorted based on the expression of cell surface marker Sca-1, we probed the identification of this protein in the *ex vivo* and *in vitro* proteomics datasets. A peptide belonging to this protein was exclusively identified in the *ex vivo* set with 7 PSMs (Fig. a). As evidenced by the relative intensity, the peptide was identified with high confidence in all the three technical replicates of the *ex vivo* sample (Fig. 4b). The inability to identify Sca-1 in the *in vitro* dataset however does not imply a lack of Sca-1 expression. Rather, the Sca-1 peptide fragmentation may be ‘masked’ by the presence of high abundance peptide species in the *in vitro* dataset as the mass spectrometry data was acquired in a data dependent acquisition (DDA) mode. This claim is validated by the analytical flow cytometry of the Sca-1^+^ cells of the *in vitro* sample, (Fig. 4c). A comparative label-free quantification of the *in vitro* Sca-1^+^ dataset was carried out with respect to the *ex vivo* set. A threshold of 1.5 fold difference in protein abundance was set to catalogue the differentially regulated proteins (Supplementary Table S3). Statistically significant proteins demonstrating variation in expression levels in *in vitro* data are represented in the volcano plot (Fig. 5a). Altogether, 1785 proteins were differentially regulated which represents approximately 30% of the proteins identified. Of the differentially regulated proteins, 1594 were upregulated and 191 were downregulated in the *in vitro* set (Fig. 5a). Among the significantly upregulated proteins, we identified cullin-associated NEDD8-dissociated protein 2 (Cand2). Interestingly, expression of Cand2 is reported to display specific expression in muscle and increased expression during myogenic differentiation [30]. This protein is reported to suppress the ubiquitination of transcription factor myogenin preventing its proteasome mediated degradation and this stabilization of myogenin promoted differentiation of C2C12 cells.

**Figure 4:**
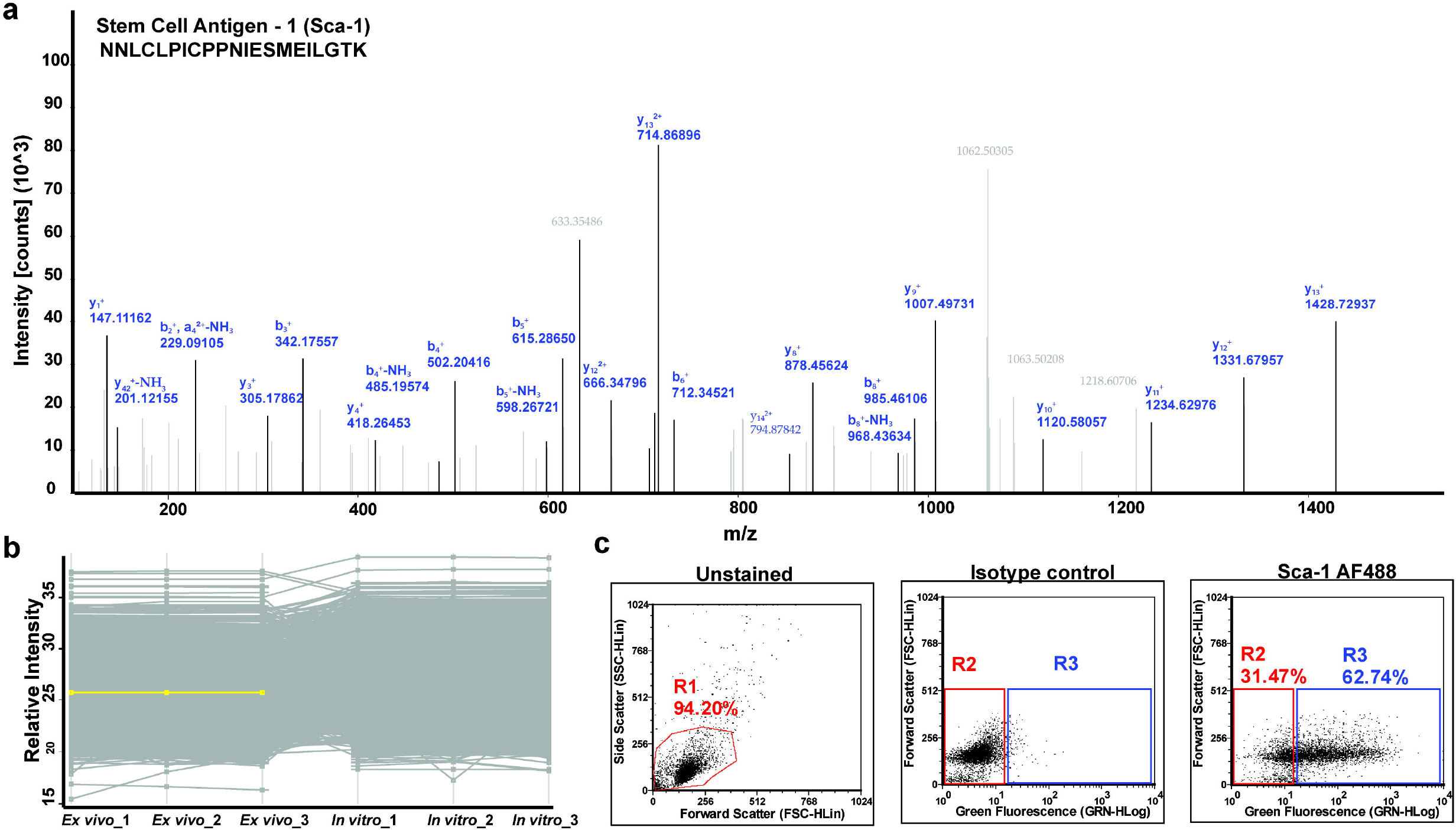
Identification of Sca-1 in mass spectrometry analysis. **(a)** Representative MS/MS spectra of Sca-1 protein identified in Sca-1^+^*ex vivo* samples. (**b**) Profile plot depicting the exclusive identification of Sca-1 in Sca-1^+^ *ex vivo* condition (Yellow line) in all three technical replicates. (**c**) Flow cytometric enumeration of Sca-1 on Sca-1^+^*in vitro* expanded cells at passage 2. Cultured cells were stained with antibody against Sca-1 for flow cytometry and analyzed using Guava^®^ easyCyte™ flow cytometer (Merck Millipore, USA). Data were analyzed using FCS Express Version 5 software.

Among the other significantly upregulated proteins, we identified dipeptidyl peptidase 3, transcriptional repressor p66-beta isoform X1, and bifunctional glutamate/proline-tRNA ligase isoform 2. Among the downregulated proteins we identified EH domain-containing protein 2, laminin subunit beta-2 precursor, histone H1.0 and myosin-4. The EH domaincontaining protein 2 (Ehd2) is reported to regulate membrane fusion of myoblast cells [31]. Unsupervised clustering of the data demonstrated global differences in protein expression patterns in the *ex vivo* and *in vitro* datasets whereas technical replicates of each set clustered together (Fig. 5b).

**Figure 5:**
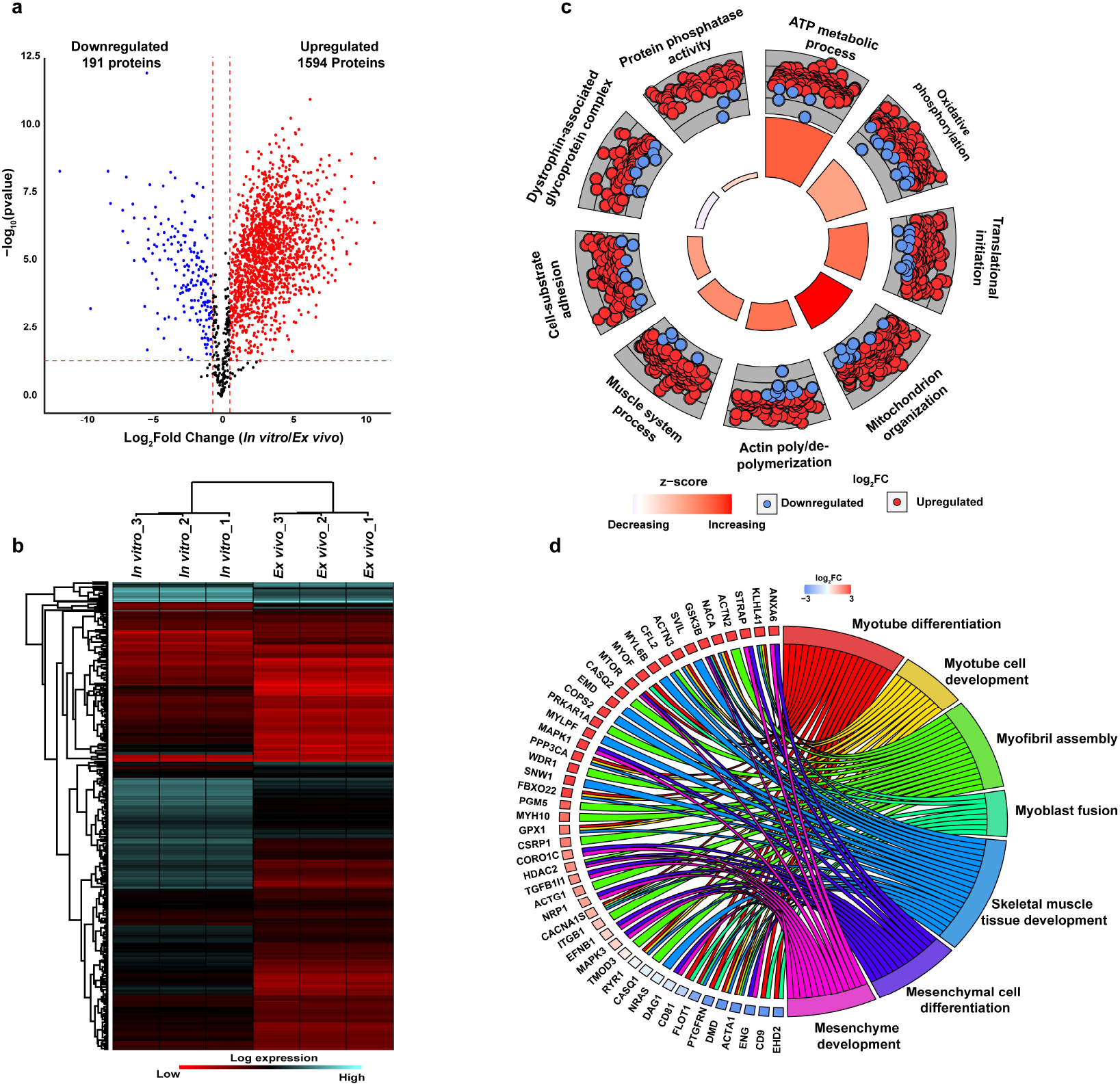
Analysis of differentially expressed proteins. (**a)** Volcano plot graph illustrating the abundance of differentially expressed proteins. The −□log10 (p value) was plotted against the log2 (ratio *in vitro /ex vivo*). **(b)** The heat map of significantly differentially expressed proteins between *ex vivo* and *in vitro* dataset. **(c)** Circular visualization (GOCircle plot) of differentially expressed proteins between *ex vivo* and *in vitro* dataset. The outer circle shows a scatter plot of differentially regulated proteins wherein the red circles indicate upregulated proteins and blue circles indicate downregulated proteins. The inner ring represents a bar plot, the height of bar represents the significance of GO term and color corresponds to z-score which measures the process is either decreased (blue) or increased (red). **(d)** GOChord plot of the GO terms involved in the skeletal muscle system processes. The gene are assigned to their respective GO terms according to their fold change.

### Biological relevance of the differentially regulated proteins

To gain a better understanding of the functional relevance of the differentially regulated proteins, GO enrichment analysis was performed. The relationship between the expression patterns of proteins in the enriched GO terms was analysed using GO Circle plot (Fig. 5c). As expected, the top enriched GO terms included metabolic processes such as ATP metabolic process and oxidative phosphorylation. Other enriched terms included translational initiation, mitochondrion organization and terms related to various aspects of muscle biology such as actin poly/depolymerisation, muscle system process, dystrophin-associated glycoprotein complex. As evident in the figure, maximum number of proteins in each GO category consisted of upregulated proteins (Fig. 5c). As several terms were enriched for muscle related functions, we examined in greater detail the representation of proteins related to regulation of muscle development and muscle regeneration in the differential proteins set (Fig. 5d). One of the proteins Annexin 6 (Anxa6) was enriched in two categories viz., mesenchymal cell differentiation and mesenchyme development. A splice variant of this protein was reported to inhibit the membrane repair in a mouse model of muscular dystrophy [32]. Protein Kelch-like protein 41 (Klhl41) was enriched in myotube differentiation, myotube cell development, myofibril assembly and skeletal muscle tissue development. Mutation in this protein has been associated with the disruption of sarcomeres leading to development of nemaline myopathy [33]. Protein serine-threonine kinase receptor-associated protein (Strap), a component of small nuclear ribonucleoproteins enriched in the categories mesenchymal cell differentiation and mesenchyme development. Previously, the overexpression of Strap led to malformations in mice [34]. As described in previous analysis, maximum proteins belonging to these categories were upregulated. As we observed a large number of kinases enriched in our differentially regulated dataset, we subjected these kinases for GO analysis to understand the pathways putatively regulated by these kinases (Fig. 6). Several pathways with relevance to muscle regeneration, myogenesis, insulin signalling, muscle atrophy, muscle stem cell maintenance and skeletal muscle maintenance were found to be enriched. The identification of several kinases among the differentially regulated proteins also hints towards the active role of protein phosphorylation in Sca-1^+^ cells under the *ex vivo* and *in vitro* conditions that can be probed in future.

**Figure 6:**
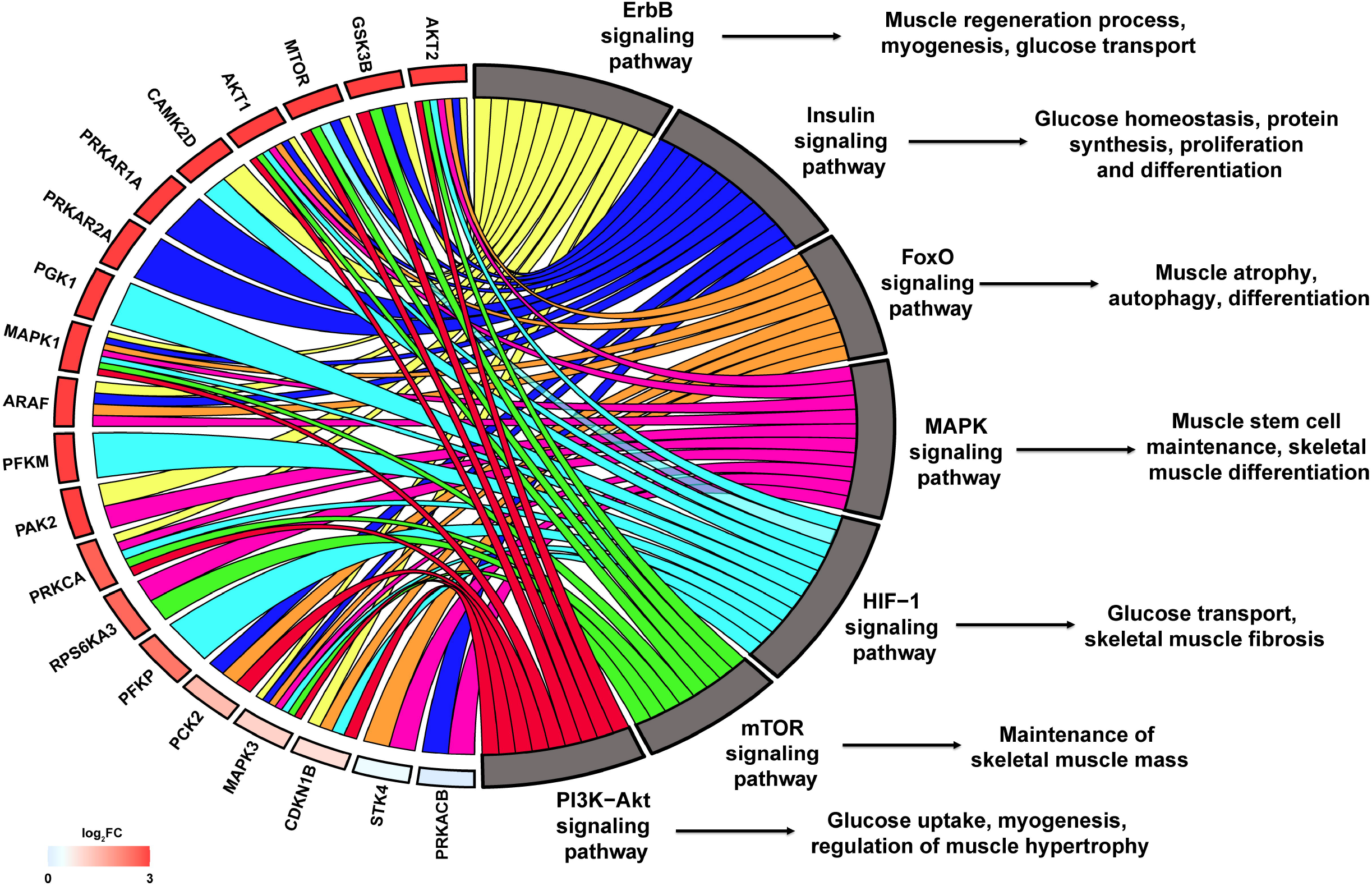
Pathways enrichment of differentially expressed proteins. GOChord plot of the GO terms involved in the various signaling pathways in skeletal muscle. The gene are assigned to their respective signaling pathway according to their fold change.

### Identification of cell surface markers

We compared all (non-redundant list of *ex vivo* and *in vitro)* the identified proteins from our study with the mouse proteins available in Cell Surface Protein Atlas (CSPA) (https://wlab.ethz.ch/cspa/#abstract). This database contains mass spectrometry-based information of mouse (and human) cell surface glycoproteins generated from 31 cell types [35]. The analysis revealed 419 proteins common to the two datasets (Supplementary Table S2). We identified several surface markers including CD177, CD44, CD47, CD59a, CD151, CD2, CD200, CD81, CD9, CD34 and Sca-1 in our dataset (Supplementary Figure S1). This also demonstrated a fair representation of membrane proteins in our data. Of the common proteins identified, 158 were found to be differentially regulated. CD44 is a well-established marker for several cancer stem cells and linked to patient prognosis [36]. However, it’s expression has been associated with the migration of myoblast as well as it’s differentiation [37]. Although, CD34 has traditionally been regarded as a marker for hematopoietic stem cells its expression on several non-hematopoietic cells including muscle satellite cells has been reported [38].

## Discussion

An overarching role of Sca-1 in skeletal muscle development and muscle regeneration is increasingly becoming evident [8,12,39,40]. Sca-1 is also a well-established marker used for the isolation of muscle derived stem cells [40]. Maintenance of stem cells under culture conditions have been reported to affect the expression of various molecules compared to their expression levels in *in vivo* state [41]. This makes it imperative to carry out the molecular profiling of freshly isolated cells and those cultured *in vitro* [41]. Here, we carried out label-free quantitative proteomics analysis to identify proteins modulated by the cellular environment in Sca-1^+^ cells. To the best of our knowledge, this is the first study on in-depth proteomic characterization of muscle derived Sca-1^+^ cells.

The deep proteomics analysis of triceps muscles and C2C12 myoblast cells represented the largest proteome dataset of mouse skeletal muscles[18]. Muscles tissues contain muscle resident adult stem cells that proliferate and give rise to various differentiated muscle cell types [42]. The low abundance of these cells makes it inevitable to speculate lack of representation of proteins expressed in low amounts or specifically expressed in these cells while carrying out tissue-based proteomics analysis. Creating deep cellular proteomics maps especially of rare cell types such as stem and progenitor cells will significantly improve our understanding of basic developmental biology.

Here, we used a FACS flow-sorting method to isolate Sca-1^+^ cells from 5-weeks-old BALB/c mice hindlimb skeletal muscle digests. We streamlined the proteomics pipeline to minimize sample loss. Starting with 100μg protein from *ex vivo* and *in vitro* datasets, we identified a total of 5581 proteins were identified. Compared to the previous study mentioned above, we identified nearly 1000 proteins unique to our study [18]. Among these proteins were Cavin1, Cavin4. These proteins are involved in formation of membrane caveolae and loss of these genes leads to metabolic disorders [43,44]. Cavin4 is reported to express exclusively in muscles [43]. Another protein exclusively identified in our study was Nucleolus and neural progenitor protein (Nepro). The role of Nepro has been characterized in development of the neocortex and hardly any information is available on its role in muscles [45]. Another protein Aff3, a transcription factor involved in neurogenesis was identified. Loss of function of Aff3 is reported to cause neurodevelopmental defects [46]. Other proteins with no known function in muscle development identified uniquely in our dataset were Nucleolar autoantigen 36 (Zfp330) and Zinc finger domain-containing protein (Fam170a). Loss or gain in function studies pertaining to such proteins without known function in muscles form the future scope of this study. The identification of unique proteins in our dataset emphasizes the importance of performing proteomics of specific population of cell types to achieve complete coverage of tissue-specific proteome.

Among the proteins uniquely identified in the *in vitro* set were a set of transcription factors including Gtf2a2, Gtf2e1, Gtf2e2, Gtf2f1, Gtf2f2, Taf10, Taf6, Tbpl1 and Mrtfb. The loss of *Taf10* in mouse embryo led to alteration of lateral plate differentiation [47]. Myocardin-related transcription factor B is reported to be reported to regulate differentiation in vascular smooth muscle cells [48]. We also identified myosin heavy chain isoforms Myh3, Myh8 and Myh15. Both Myh3 and Myh8 are reported to express specifically in the embryonic and fetal stages following which they are only re-expressed during muscle regeneration [49]. Expression of Taf6 is reported in the TFIID complex of myoblasts and differentiated myotubes [50]. A large panel of spliceosome associated factors, mitochondrial ribosomal proteins, ankyrin repeat proteins and RNA transport related molecules were also identified exclusively in the *in vitro* set.

We identified marker proteins that are known to express on muscle cell types. High confidence identification of endogenous Sca-1 was observed exclusively in the *ex vivo* dataset. However, we confirmed the expression of Sca-1 in the *in vitro* sample using analytical flow cytometry. Possibly, the identification of Sca-1 in the *in vitro* sample mass spectrometry data may have been masked by the relatively more abundant proteins. Sca-1 has been reported to negatively regulate the proliferation and differentiation of myoblasts [8]. The lack of Sca-1 expression in mice led to hyperproliferation of myoblasts in response to injury and delay in muscle repair [51]. We can speculate here that the Sca1^+^ cells in the mouse hindlimb muscles may be indicative of homeostasis of muscle repair and regulation and proliferation of myoblasts. Among other cell surface markers identified, CD34 has principally been used as a marker for HSCs. However, its expression on multiple other cell types has been documented [38]. CD34 has been reported as a novel marker expressed on quiescent muscle satellite cells [52]. The identification of CD34 in our data warrants further investigation to determine its exact role in muscle development. Cell surface markers such as the CD34, CD44, CD24 and CD15 are regulated through extensive glycosylation. In future, it will be interesting to probe the roles of glycosylation in regulating stem/progenitor cell properties.

We also identified a panel of upregulated kinases including Mtor, Akt1, Akt2, Mapk1, Mapk3, Cdk9, Rock1, Camk2d and Rack1. Depletion of mTOR complex mTORC1 led to defects in myogenesis and death in mice [53]. Conditional knockout of *mTOR* exhibited severe defects in muscle regeneration and reduced size of regenerated myofibers and the myoblasts expressed lower levels of myogenic determinant genes [54]. mTOR lies downstream of phosphatidylinositol 3-kinase (PI3K)/AKT and Ras/MAPK signalling pathways that together control protein synthesis [55]. Phosphorylation of TSC2 by Akt leads to a cascade of molecular events that activate mTOR which in turn phosphorylates eukaryotic translation initiation factor 4E-binding protein (Eif4ebp1) [56]. We also identified Eif4ebp1 as an upregulated protein indicating increased protein translation under *in vitro* condition.

Among other proteins related to translation, we identified components of the 40S ribosomal protein (Rps2-Rps28) and several eukaryotic translation initiation factors and their binding proteins. Expression of Akt is associated with induction of hypertrophy in skeletal muscles [57]. Akt has also been reported to maintain metabolic homeostasis in muscles, and loss of Akt2 led to glucose intolerance and insulin resistant phenotype [58]. Insulin-like growth factors (Igfs) are also associated with hypertrophy [59]. Igf1 is positively associated with regulation of the size of myofibres, a process that also involves the phosphorylation of Akt [59]. We identified insulin-like growth factor-binding protein 2 isoform 1 as an upregulated protein. In aged mice, decreased regenerative capacity of satellite cells has been associated with increase in Igfbp-2 and Igfbp-3 and decrease in insulin-like growth factor-II [60]. We also identified two isoforms of glycogen synthase kinase-3 (Gsk3a and Gsk3b), both of which were upregulated. Gsk3a has been reported to regulate the activity of mTORC1 [61]. Our data support the assumption that under *in vitro* condition, Sca-1^+^ cells display increased protein synthesis. The *in vitro* environment also triggered hypertrophy that could have resulted through the activation of muscle Sca-1^+^ cells [62]. A partial list of protein identified in present study is presented in Table 1.

**Table 1:** Partial list of proteins identified in the present study

| **Protein** | **Expression in our data** | **Known function in muscle development/protein translation/stem cell regulation** | **Reference** |
| --- | --- | --- | --- |
| **Cand2** | Upregulated in *in vitro* | Myogenic differentiation | 30 |
| **mTOR** | Upregulated in *in vitro* | Control of protein synthesis; muscle regeneration | 54 |
| **4E-BP1** | Upregulated in *in vitro* | Inhibitor of translational initiation | 56 |
| **Akt2** | Upregulated in *in vitro* | Loss of gene led to glucose intolerance and insulin resistant phenotype | 58 |
| **Igfbp2** | Upregulated in *in vitro* | Muscle regeneration | 60 |
| **Gsk3a** | Upregulated in *in vitro* | Regulate the activity of mTORC1 | 61 |
| **STAT3** | Upregulated in *in vitro* | Muscle hypertrophy; regulation of MyoD1 | 75 |
| **CD44** | Upregulated in *in vitro* | Migration of myoblasts and differentiation | 37 |
| **Mrtfb** | Exclusive to *in vitro* | Differentiation in vascular smooth muscle cells | 48 |
| **CAVIN4** | Exclusive to *in vitro* | Expression restricted to striated muscles | 43 |
| **Sca-1** | Exclusive to ex vivo | Muscle development and muscle regeneration | 8, 12, 39, 40 |
| **Stk10** | Downregulated in *in vitro* | No known function | NA |
| **Anxa6** | Upregulated in *in vitro* | Muscle membrane leak and repair | 32 |
| **EHD2** | Downregulated in *in vitro* | Membrane fusion of myoblast cells | 31 |

Among the downregulated proteins, we identified one of the core components of the Hippo signalling pathway, protein serine/threonine-protein kinase 10 isoform 1. Loss of function of the Drosophila pro-apoptotic kinase *hippo* (Stk3/Mst2 and Stk4/Mst1, in mammals) resulted in enlarged organs [63]. Largely, Mst1/2 have been characterized as negative regulators of hyperproliferation and tumorigenesis [64]. However, mice deficient for both these kinases died *in utero* indicating their role in developmental biology [65]. However, there is hardly any information on Stk10 with respect to muscle development.

We also identified transcription factors in our dataset. Sp1 transcription factor was reported to activate Myod and promote muscle cell differentiation [66]. In our data, Sp1 was exclusively identified in the *in vitro* set. Another upregulated protein that was enriched in multiple pathways including translation was Stat3. Proliferating satellite cells activate Stat3 signalling pathways and contribute to hypertrophy [67]. Activation of this protein is regulated through phosphorylation by multiple kinases including Mtor kinase that increases its transcriptional activity [68]. In muscle satellite cells, Stat3 promotes myogenic lineage progression through regulation of Myod1 [69]. Overall, our results reveal that under *in vitro* condition, Sca-1^+^ cells display increased protein synthesis. This is reflected by increased expression of Mtor, its well-characterized substrate Eif4ebp1 and multiple subunits of ribosomal protein. An important segment of our analysis is also the identification of several cell surface markers that in future may be used as surrogate molecules to isolate Sca-1^+^ cells. Designing such combinatorial strategies for the isolation of Sca-1^+^ cells will provide higher validity to enrichment scores for isolation of pure cell populations.

## Acknowledgements

The authors would like to thank the Stem Cells and Regenerative Medicine Centre of Yenepoya Research Centre, Yenepoya (Deemed to be University) for providing the infrastructure, core facility and funding in the form of Yenepoya University Seed Grant (YU/Seed Grant/2015-042) awarded to the Principal Investigator Dr Bipasha Bose to carry out this study. The authors would like to acknowledge the MASSFIITB Facility at IIT Bombay supported by the Department of Biotechnology (BT/PR13114/INF/22/206/2015) to carry out all MS-related experiments.

## Author contributions

BB, SK and SS conceived and designed the study. SK and SS performed the animal experiments. SK carried out the MS sample preparation, analysed and performed the data analysis. SK and PS wrote the manuscript. BB and SS edited and approved the manuscript.

## Supplementary files

**Table S1:** The entire list of proteins identified in the Sca-1^+^ *ex vivo* and *in vitro* conditions.

**Table S2:** List of regulatory proteins identified in this study.

**Table S3:** List of proteins which are differentially expressed between Sca-1^+^*ex vivo* and *in vitro* condition.

## References

1. Epting, C. L., King, F. W., Pedersen, A., et al. (2008). Stem cell antigen-1 localizes to lipid microdomains and associates with insulin degrading enzyme in skeletal myoblasts. J Cell Physiol, 217, 250–260.

2. Spangrude, G. J., Heimfeld, S., Weissman, I. L. (1988). Purification and characterization of mouse hematopoietic stem cells. Science, 241, 58–62.

3. Holmes, C., Stanford, W. L. (2007). Concise review: stem cell antigen-1: expression, function, and enigma. Stem Cells, 25, 1339–1347.

4. Lee, J. Y., Qu-Petersen, Z., Cao, B., et al. (2000). Clonal isolation of muscle-derived cells capable of enhancing muscle regeneration and bone healing. J Cell Biol, 150, 1085–1100.

5. Shen, X., Collier, J. M., Hlaing, M., et al. (2003). Genome-wide examination of myoblast cell cycle withdrawal during differentiation. Dev Dyn, 226, 128–138.

6. Zammit, P., Beauchamp, J. (2001). The skeletal muscle satellite cell: stem cell or son of stem cell? Differentiation, 68, 193–204.

7. Judson, R. N., Low, M., Eisner, C., Rossi, F. M. (2017). Isolation, Culture, and Differentiation of Fibro/Adipogenic Progenitors (FAPs) from Skeletal Muscle. Methods Mol Biol, 1668, 93–103.

8. Mitchell, P. O., Mills, T., O’Connor, R. S., et al. (2005). Sca-1 negatively regulates proliferation and differentiation of muscle cells. Dev Biol, 283, 240–252.

9. Torrente, Y., Tremblay, J. P., Pisati, F., et al. (2001). Intraarterial injection of muscle-derived CD34(+)Sca-1(+) stem cells restores dystrophin in mdx mice. J Cell Biol, 152, 335–348.

10. Oh, H., Bradfute, S. B., Gallardo, T. D., et al. (2003). Cardiac progenitor cells from adult myocardium: homing, differentiation, and fusion after infarction. Proc Natl Acad Sci U S A, 100, 12313–12318.

11. Bonyadi, M., Waldman, S. D., Liu, D., et al. (2003). Mesenchymal progenitor self-renewal deficiency leads to age-dependent osteoporosis in Sca-1/Ly-6A null mice. Proc Natl Acad Sci US A, 100, 5840–5845.

12. Kafadar, K. A., Yi, L., Ahmad, Y., et al. (2009). Sca-1 expression is required for efficient remodeling of the extracellular matrix during skeletal muscle regeneration. Dev Biol, 326, 47–59.

13. Epting, C. L., Lopez, J. E., Shen, X., et al. (2004). Stem cell antigen-1 is necessary for cell-cycle withdrawal and myoblast differentiation in C2C12 cells. J Cell Sci, 117, 6185–6195.

14. Swietlik, J. J., Sinha, A., Meissner, F. (2020). Dissecting intercellular signaling with mass spectrometry-based proteomics. Curr Opin Cell Biol, 63, 20–30.

15. Gstaiger, M., Aebersold, R. (2009). Applying mass spectrometry-based proteomics to genetics, genomics and network biology. Nat Rev Genet, 10, 617–627.

16. Amon, S., Meier-Abt, F., Gillet, L. C., et al. (2019). Sensitive Quantitative Proteomics of Human Hematopoietic Stem and Progenitor Cells by Data-independent Acquisition Mass Spectrometry. Mol Cell Proteomics, 18, 1454–1467.

17. Ohlendieck, K. (2011). Skeletal muscle proteomics: current approaches, technical challenges and emerging techniques. Skelet Muscle, 1, 6.

18. Deshmukh, A. S., Murgia, M., Nagaraj, N., et al. (2015). Deep proteomics of mouse skeletal muscle enables quantitation of protein isoforms, metabolic pathways, and transcription factors. Mol Cell Proteomics, 14, 841–853.

19. Kleinert, M., Parker, B. L., Jensen, T. E., et al. (2018). Quantitative proteomic characterization of cellular pathways associated with altered insulin sensitivity in skeletal muscle following high-fat diet feeding and exercise training. Sci Rep, 8, 10723.

20. Ubaida-Mohien, C., Gonzalez-Freire, M., Lyashkov, A., et al. (2019). Physical Activity Associated Proteomics of Skeletal Muscle: Being Physically Active in Daily Life May Protect Skeletal Muscle From Aging. Front Physiol, 10, 312.

21. Yin, X., Mayr, M., Xiao, Q., et al. (2005). Proteomic dataset of Sca-1+ progenitor cells. Proteomics, 5, 4533–4545.

22. Sudheer Shenoy, P., Bose, B. (2017). Identification, isolation, quantification and systems approach towards CD34, a biomarker present in the progenitor/stem cells from diverse lineages. Methods, 131, 147–156.

23. Kulak, N. A., Pichler, G., Paron, I., Nagaraj, N., Mann, M. (2014). Minimal, encapsulated proteomic-sample processing applied to copy-number estimation in eukaryotic cells. Nat Methods, 11, 319–324.

24. Walter, W., Sanchez-Cabo, F., Ricote, M. (2015). GOplot: an R package for visually combining expression data with functional analysis. Bioinformatics, 31, 2912–2914.

25. Tapscott, S. J. (2005). The circuitry of a master switch: Myod and the regulation of skeletal muscle gene transcription. Development, 132, 2685–2695.

26. Molkentin, J. D., Olson, E. N. (1996). Combinatorial control of muscle development by basic helix-loop-helix and MADS-box transcription factors. Proc Natl Acad Sci U S A, 93, 9366–9373.

27. Sartorelli, V., Caretti, G. (2005). Mechanisms underlying the transcriptional regulation of skeletal myogenesis. Curr Opin Genet Dev, 15, 528–535.

28. Rotwein, P., Wilson, E. M. (2009). Distinct actions of Akt1 and Akt2 in skeletal muscle differentiation. J Cell Physiol, 219, 503–511.

29. Forrest, A. R., Ravasi, T., Taylor, D., et al. (2003). Phosphoregulators: protein kinases and protein phosphatases of mouse. Genome Res, 13, 1443–1454.

30. Shiraishi, S., Zhou, C., Aoki, T., et al. (2007). TBP-interacting protein 120B (TIP120B)/cullin-associated and neddylation-dissociated 2 (CAND2) inhibits SCF-dependent ubiquitination of myogenin and accelerates myogenic differentiation. J Biol Chem, 282, 9017–9028.

31. Doherty, K. R., Demonbreun, A. R., Wallace, G. Q., et al. (2008). The endocytic recycling protein EHD2 interacts with myoferlin to regulate myoblast fusion. J Biol Chem, 283, 20252–20260.

32. Swaggart, K. A., Demonbreun, A. R., Vo, A. H., et al. (2014). Annexin A6 modifies muscular dystrophy by mediating sarcolemmal repair. Proc Natl Acad Sci U S A, 111, 6004–6009.

33. Ramirez-Martinez, A., Cenik, B. K., Bezprozvannaya, S., et al. (2017). KLHL41 stabilizes skeletal muscle sarcomeres by nonproteolytic ubiquitination. Elife, 6.

34. Zhang, X., Azhar, G., Rogers, S. C., et al. (2014). Overexpression of p49/STRAP alters cellular cytoskeletal structure and gross anatomy in mice. BMC Cell Biol, 15, 32.

35. Bausch-Fluck, D., Hofmann, A., Bock, T., et al. (2015). A mass spectrometric-derived cell surface protein atlas. PLoS One, 10, e0121314.

36. Senbanjo, L. T., Chellaiah, M. A. (2017). CD44: A Multifunctional Cell Surface Adhesion Receptor Is a Regulator of Progression and Metastasis of Cancer Cells. Front Cell Dev Biol, 5, 18.

37. Mylona, E., Jones, K. A., Mills, S. T., Pavlath, G. K. (2006). CD44 regulates myoblast migration and differentiation. J Cell Physiol, 209, 314–321.

38. Sidney, L. E., Branch, M. J., Dunphy, S. E., Dua, H. S., Hopkinson, A. (2014). Concise review: evidence for CD34 as a common marker for diverse progenitors. Stem Cells, 32, 1380–1389.

39. Mann, C. J., Perdiguero, E., Kharraz, Y., et al. (2011). Aberrant repair and fibrosis development in skeletal muscle. Skelet Muscle, 1, 21.

40. Bernstein, H. S., Samad, T., Cholsiripunlert, S., et al. (2013). Stem cell antigen-1 in skeletal muscle function. PLoS Curr, 5.

41. Kim, T., Echeagaray, O. H., Wang, B. J., et al. (2018). In situ transcriptome characteristics are lost following culture adaptation of adult cardiac stem cells. Sci Rep, 8, 12060.

42. Wang, Y. X., Dumont, N. A., Rudnicki, M. A. (2014). Muscle stem cells at a glance. J Cell Sci, 127, 4543–4548.

43. Liu, L., Hansen, C. G., Honeyman, B. J., Nichols, B. J., Pilch, P. F. (2014). Cavin-3 knockout mice show that cavin-3 is not essential for caveolae formation, for maintenance of body composition, or for glucose tolerance. PLoS One, 9, e102935.

44. Liu, L., Brown, D., McKee, M., et al. (2008). Deletion of Cavin/PTRF causes global loss of caveolae, dyslipidemia, and glucose intolerance. Cell Metab, 8, 310–317.

45. Saito, T. (2012). NEPRO: a novel Notch effector for maintenance of neural progenitor cells in the neocortex. Adv Exp Med Biol, 727, 61–70.

46. Moore, J. M., Oliver, P. L., Finelli, M. J., et al. (2014). Laf4/Aff3, a gene involved in intellectual disability, is required for cellular migration in the mouse cerebral cortex. PLoS One, 9, e105933.

47. Bardot, P., Vincent, S. D., Fournier, M., et al. (2017). The TAF10-containing TFIID and SAGA transcriptional complexes are dispensable for early somitogenesis in the mouse embryo. Development, 144, 3808–3818.

48. Xie, W. B., Li, Z., Shi, N., et al. (2013). Smad2 and myocardin-related transcription factor B cooperatively regulate vascular smooth muscle differentiation from neural crest cells. Circ Res, 113, e76–86.

49. Schiaffino, S., Rossi, A. C., Smerdu, V., Leinwand, L. A., Reggiani, C. (2015). Developmental myosins: expression patterns and functional significance. Skelet Muscle, 5, 22.

50. Malecova, B., Dall’Agnese, A., Madaro, L., et al. (2016). TBP/TFIID-dependent activation of MyoD target genes in skeletal muscle cells. Elife, 5.

51. Epting, C. L., Lopez, J. E., Pedersen, A., et al. (2008). Stem cell antigen-1 regulates the tempo of muscle repair through effects on proliferation of alpha7 integrin-expressing myoblasts. Exp Cell Res, 314, 1125–1135.

52. Beauchamp, J. R., Heslop, L., Yu, D. S., et al. (2000). Expression of CD34 and Myf5 defines the majority of quiescent adult skeletal muscle satellite cells. J Cell Biol, 151, 1221–1234.

53. Rion, N., Castets, P., Lin, S., et al. (2019). mTOR controls embryonic and adult myogenesis via mTORC1. Development, 146.

54. Zhang, P., Liang, X., Shan, T., et al. (2015). mTOR is necessary for proper satellite cell activity and skeletal muscle regeneration. Biochem Biophys Res Commun, 463, 102–108.

55. Kim, H. J. (2019). Cell Fate Control by Translation: mRNA Translation Initiation as a Therapeutic Target for Cancer Development and Stem Cell Fate Control. Biomolecules, 9.

56. Inoki, K., Li, Y., Zhu, T., Wu, J., Guan, K. L. (2002). TSC2 is phosphorylated and inhibited by Akt and suppresses mTOR signalling. Nat Cell Biol, 4, 648–657.

57. Lai, K. M., Gonzalez, M., Poueymirou, W. T., et al. (2004). Conditional activation of akt in adult skeletal muscle induces rapid hypertrophy. Mol Cell Biol, 24, 9295–9304.

58. Jaiswal, N., Gavin, M. G., Quinn, W. J., 3rd, et al. (2019). The role of skeletal muscle Akt in the regulation of muscle mass and glucose homeostasis. Mol Metab, 28, 1–13.

59. Hennebry, A., Oldham, J., Shavlakadze, T., et al. (2017). IGF1 stimulates greater muscle hypertrophy in the absence of myostatin in male mice. J Endocrinol, 234, 187–200.

60. Ikemoto-Uezumi, M., Uezumi, A., Tsuchida, K., et al. (2015). Pro-Insulin-Like Growth Factor-II Ameliorates Age-Related Inefficient Regenerative Response by Orchestrating Self-Reinforcement Mechanism of Muscle Regeneration. Stem Cells, 33, 2456–2468.

61. Zhou, J., Freeman, T. A., Ahmad, F., et al. (2013). GSK-3alpha is a central regulator of age-related pathologies in mice. J Clin Invest, 123, 1821–1832.

62. Fukada, S. I., Akimoto, T., Sotiropoulos, A. (2020). Role of damage and management in muscle hypertrophy: Different behaviors of muscle stem cells in regeneration and hypertrophy. Biochim Biophys Acta Mol Cell Res, 1867, 118742.

63. Wu, S., Huang, J., Dong, J., Pan, D. (2003). hippo encodes a Ste-20 family protein kinase that restricts cell proliferation and promotes apoptosis in conjunction with salvador and warts. Cell, 114, 445–456.

64. Qin, F., Tian, J., Zhou, D., Chen, L. (2013). Mst1 and Mst2 kinases: regulations and diseases. Cell Biosci, 3, 31.

65. Oh, S., Lee, D., Kim, T., et al. (2009). Crucial role for Mst1 and Mst2 kinases in early embryonic development of the mouse. Mol Cell Biol, 29, 6309–6320.

66. Guo, C. S., Degnin, C., Fiddler, T. A., Stauffer, D., Thayer, M. J. (2003). Regulation of MyoD activity and muscle cell differentiation by MDM2, pRb, and Sp1. J Biol Chem, 278, 22615–22622.

67. Serrano, A. L., Baeza-Raja, B., Perdiguero, E., Jardi, M., Munoz-Canoves, P. (2008). Interleukin-6 is an essential regulator of satellite cell-mediated skeletal muscle hypertrophy. Cell Metab, 7, 33–44.

68. Sala, D., Sacco, A. (2016). Signal transducer and activator of transcription 3 signaling as a potential target to treat muscle wasting diseases. Curr Opin Clin Nutr Metab Care, 19, 171–176.

69. Tierney, M. T., Aydogdu, T., Sala, D., et al. (2014). STAT3 signaling controls satellite cell expansion and skeletal muscle repair. Nat Med, 20, 1182–1186.

